# ERGA-BGE Reference Genome of *Phyllidia flava* (Nudibranchia: Phyllidiidae), a Relict Species Endemic to the Mediterranean Sea

**DOI:** 10.1101/2025.01.14.632929

**Authors:** Carles Galià-Camps, Miquel Pontes, Thomas Marcussen, Torsten H. Struck, Rebekah Oomen, Astrid Böhne, Rita Monteiro, Laura Aguilera, Marta Gut, Francisco Câmara Ferreira, Fernando Cruz, Jèssica Gómez-Garrido, Tyler S. Alioto, Diego De Panis

## Abstract

*Phyllidia flava*, the only Mediterranean representative of the otherwise Indo-Pacific family Phyllididae, is a radula-less dorid nudibranch that feeds exclusively on toxic sponges, from which it derives defensive and camouflage compounds. Although not currently listed as endangered, this relict species remains vulnerable due to its restricted distribution and the rising temperatures of the Mediterranean Sea. A total of 13 contiguous chromosomal pseudomolecules were assembled, resulting in a chromosome-level genome spanning 1.9 Gb. This assembly comprises 2,032 contigs and 62 scaffolds, with contig and scaffold N50 values of 2.4 Mb and 169.7 Mb, respectively. This reference genome will serve as a valuable resource for conservation efforts, provide a robust calibration point for studying nudibranch phylogenetics based on the Tethys Ocean closure, and lay the groundwork for research into toxic prey diet specialisation.

## Introduction

*Phyllidia flava* Aradas, 1847 is the only Phyllididae species endemic to the Mediterranean Sea, with all other members of the family living in the Indian and Pacific Oceans. Adult *P. flava* specimens reach sizes up to five centimeters and are characterised by an orange-to-yellow body with small whitish warts on their mantle. Like all the members of the superfamily Phyllidioidea, it is a radula-less dorid species (Galià-Camps et al., 2022, 2024; Papu et al., 2022), being the radula one of the most relevant taxonomic traits for nudibranch identification. Additionally, Phyllididae species possess a pleural gill, a rare trait among nudibranchs, which allows easy identification from species belonging to related families (Brunckhorst, 1993). *Phyllidia flava* inhabits two scarce toxic sponges, *Axinella cannabina* and *Acanthella acuta*, the latter being its primary food source (Furfaro & Mariottini, 2016; Wägele & Klussmann-Kolb, 2005). Remarkably, P. flava is one of the few known species to prey on these Mediterranean sponges, playing a vital role in regulating their populations and maintaining ecosystem balance. Through predation, *P. flava* acquires the necessary compounds to metabolise deterrent sesquiterpenes as its main chemical defense mechanism (Carbone et al., 2019) and incorporates siliceous spicules to strengthen its mantle, mimicking the sponge’s texture for camouflage (Wägele & Klussmann-Kolb, 2005).

Although not currently listed as endangered by the IUCN, *P. flava* is highly vulnerable due to its restricted distribution, small population sizes, and the alarming warming of Mediterranean waters.

Developing a high-quality reference genome for *Phyllidia flava* will significantly advance research across multiple disciplines, especially ecology and evolution. As the sole representative species of *Phyllidia* in the Mediterranean, this genome will serve as a critical calibration point in nudibranch and mollusc phylogenetic studies, tracing the divergence from its sibling species following the closure of the Tethys Sea, approximately 20 million years ago (Torfstein & Steinberg, 2020). This genome will provide valuable insights into the evolution of molluscan synapomorphies, such as shell and radula formation, while shedding light on molecular mechanisms underlying strict prey-diet relationships. These insights will deepen our understanding of ecological interactions. Furthermore, this genomic resource will facilitate genetic population assessments, aiding in the evaluation of *P. flava’s* conservation status. This could support its inclusion on the IUCN red list and bolster its role as a flagship species, raising public awareness of the Mediterranean ecosystem’s uniqueness and the need for its conservation.

The generation of this reference resource was coordinated by the European Reference Genome Atlas (ERGA) initiative’s Biodiversity Genomics Europe (BGE) project, supporting ERGA’s aims of promoting transnational cooperation to promote advances in the application of genomics technologies to protect and restore biodiversity (Mazzoni et al., 2023). This species falls within the regional reach of the Catalan Initiative for the Earth BioGenome Project (CBP) (Corominas et al., 2024), which is affiliated with ERGA.

## Materials & Methods

ERGA’s sequencing strategy includes Oxford Nanopore Technology (ONT) and/or Pacific Biosciences (PacBio) for long-read sequencing, along with Hi-C sequencing for chromosomal architecture, Illumina Paired-End (PE) for polishing (i.e. recommended for ONT-only assemblies), and RNA sequencing for transcriptomic profiling, to facilitate genome assembly and annotation.

### Sample and Sampling Information

On July 28^th^, 2023, an adult specimen of *Phyllidia flava* was sampled and photographed by Miquel Pontes. The species was initially identified in the field based on its morphological traits and unique ecological association with *Acanthella acuta* (being the only *Phyllidia* species in the Mediterranean Sea that lives on this sponge). This identification was later confirmed using a local nudibranch field guide (Ballesteros et al., 2019), under the coordination of Carles Galià-Camps. The specimen was manually collected from a sponge during a scuba dive at a depth of 25 m near Es Caials in Cadaqués, Girona (Spain). Sampling was conducted under permits issued by the Catalan Government’s Department of Climate Action, Food, and Rural Agenda (permit number DG051201-333/2022). The specimen’s tissues (mollusc foot and hepatopancreas, among others) were independently snap-frozen immediately after harvesting and stored in liquid nitrogen until DNA extraction.

### Vouchering information

Physical reference materials for the sequenced specimen (Figure 1) have been deposited in the National Museum of Natural Sciences of Madrid (Spain), under the accession number MNCN15.06/96127. An electronic voucher image of the sequenced individual is available from ERGA’s EBI BioImageArchive dataset www.ebi.ac.uk/biostudies/bioimages/studies/S-BIAD1012?query=ERGA under accession ID SAMEA114519163.

**Figure 1.**
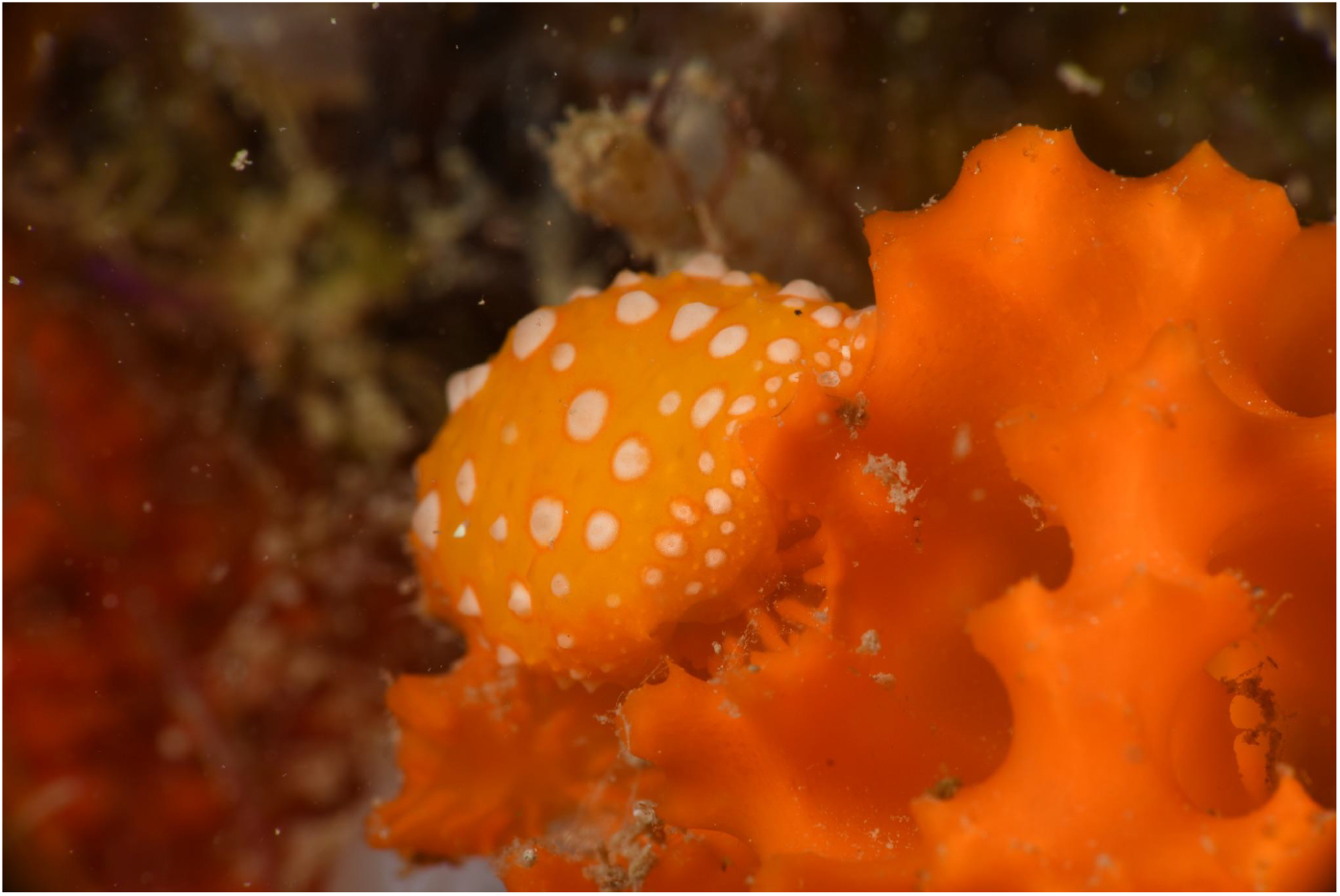
The sequenced individual of *Phyllidia flava* feeding on *Acanthella acuta*. Photograph by Miquel Pontes. This image (SAMEA114519163_5.JPG), along with five others, is part of the electronic voucher available in ERGA’s EBI BioImageArchive dataset.

### Data Availability

*Phyllidia flava* and the related genomic study were assigned to Tree of Life ID (ToLID) ‘xgPhyFlav1’ and all sample, sequence, and assembly information are available under the umbrella BioProject PRJEB77892. The sample information is available at the following BioSample accessions: SAMEA114519166, SAMEA114519168, SAMEA114519172, and SAMEA114519174. The genome assembly is accessible from ENA under accession number GCA_964266695.1. Sequencing data produced as part of this project are available from ENA at the following accessions: ERX12756248, ERX12756249, ERX13167082, and ERX13167083. Documentation related to the genome assembly and curation can be found in the ERGA Assembly Report (EAR) available at github.com/ERGA-consortium/EARs/tree/main/Assembly_Reports/Phyllidia_flava/xgPhyFlav1.

Further details and data about the project are hosted on the ERGA portal at portal.erga-biodiversity.eu/data_portal/2934182.

### Genetic Information

The estimated genome size, based on ancestral taxa, is 1.96 Gb, while the estimation based on reads kmer profiling is 2.24 Gb. This is a diploid hermaphrodite and monoecious species with an estimated haploid number based on ancestral taxa of 12 chromosomes (2n=24). Information for this species was retrieved from Genomes on a Tree (Challis et al., 2023).

### DNA processing

DNA was extracted from the foot and mantle using the Blood & Cell Culture DNA Mini Kit (Qiagen) following the manufacturer’s instructions. DNA quantification was performed using a Qubit dsDNA BR Assay Kit (Thermo Fisher Scientific), and DNA integrity was assessed using a Femtopulse system (Genomic DNA 165 Kb Kit, Agilent). DNA was stored at 4^º^C until use.

### Library Preparation and Sequencing

For long-read whole genome sequencing (WGS), a library was prepared using the SQK-LSK114 kit and sequenced on a PromethION P24 A series instrument (Oxford Nanopore Technologies). For short-read WGS, a library was prepared using the KAPA Hyper Prep Kit (Roche). A Hi-C library was prepared from hepatopancreas using the High Coverage Hi-C Kit (ARIMA), followed by the KAPA Hyper Prep Kit for Illumina sequencing (Roche). All the short-read libraries were sequenced on the Illumina NovaSeq 6000 instrument. In total, 72x Oxford Nanopore, 74x Illumina WGS shotgun, and 64x Hi-C data were sequenced to generate the assembly.

### Genome Assembly Methods

The genome was assembled using the CNAG CLAWS pipeline (Gomez-Garrido, 2024). Briefly, reads were preprocessed for quality and length using Trim Galore v0.6.7 and Filtlong v0.2.1, and initial contigs were assembled using NextDenovo v2.5.0, followed by polishing of the assembled contigs using HyPo v1.0.3, removal of retained haplotigs using purge-dups v1.2.6 and scaffolding with YaHS v1.2a. Finally, assembled scaffolds were curated via manual inspection using Pretext v0.2.5 with the Rapid Curation Toolkit (gitlab.com/wtsi-grit/rapid-curation) to remove any false joins and incorporate any sequences not automatically scaffolded into their respective locations in the chromosomal pseudomolecules (or super-scaffolds). Summary analysis of the released assembly was performed using the ERGA-BGE Genome Report ASM Galaxy workflow (De Panis, 2024), incorporating tools such as BUSCO v5.5, Merqury v1.3, and others (see reference for the full list of tools).

## Results

### Genome Assembly

The genome assembly has a total length of 1,900,507,057 bp in 62 scaffolds (Figures 2 and 3), with a GC content of 36.18%. It features a contig N50 of 2,379,035 bp (L50=214) and a scaffold N50 of 169,704,197 bp (L50=4). There are 1,970 gaps, totaling 394,000 bp in cumulative size. The single-copy gene content analysis using the metazoa database with BUSCO resulted in 95.6% completeness (95.2% single and 0.4% duplicated). 74.45% of reads k-mers were present in the assembly and the assembly has a base accuracy Quality Value (QV) of 40.42 as calculated by Merqury.

**Figure 2.**
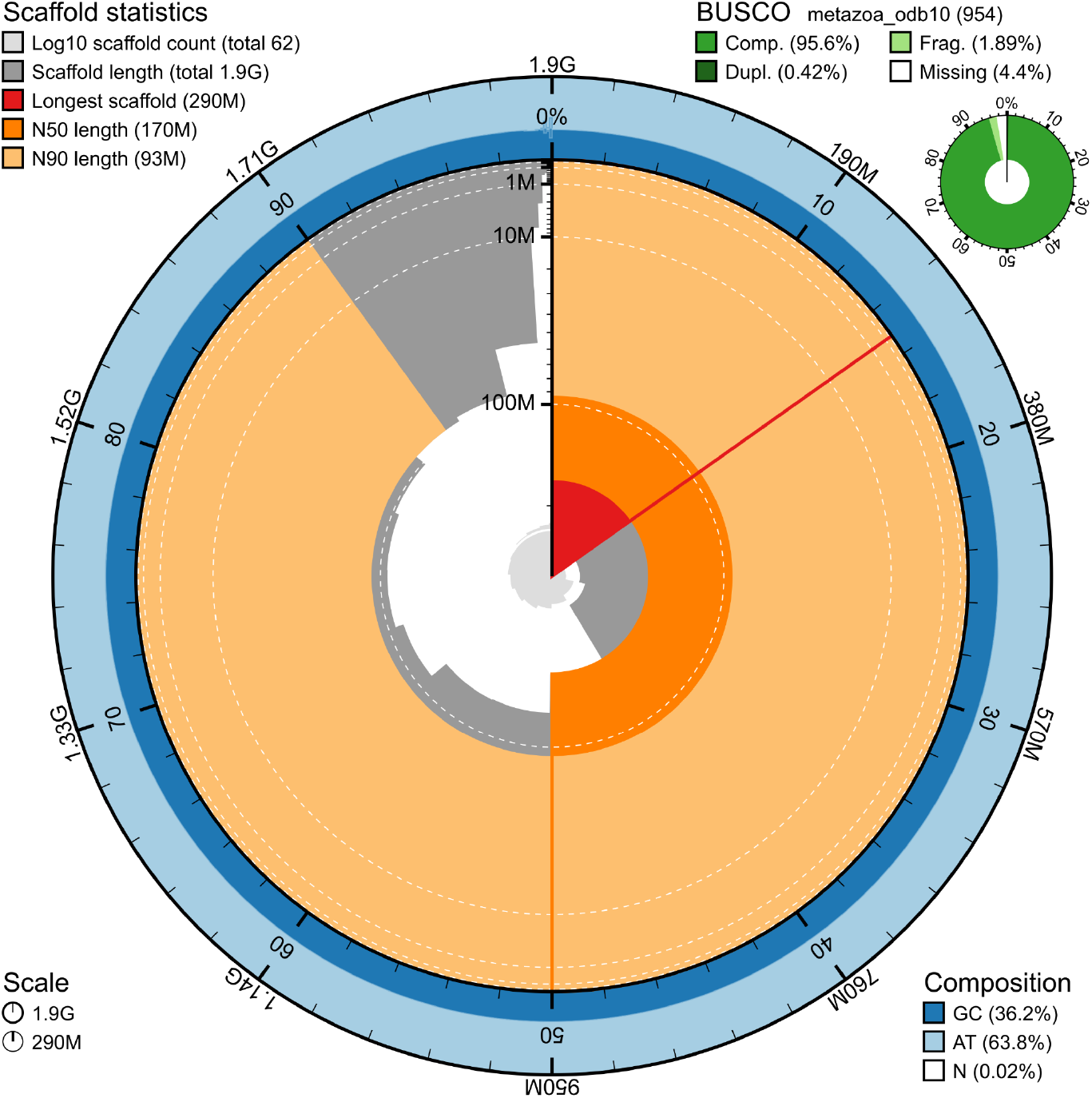
Snail plot summary of assembly statistics. The main plot is divided into 1,000 size-ordered bins around the circumference, with each bin representing 0.1% of the 1,900,507,057 bp assembly. The distribution of sequence lengths is shown in dark grey, with the plot radius scaled to the longest sequence present in the assembly (289,735,081 bp, shown in red). Orange and pale-orange arcs show the scaffold N50 and N90 sequence lengths (169,704,197 and 92,966,410 bp), respectively. The pale grey spiral shows the cumulative sequence count on a log-scale, with white scale lines showing successive orders of magnitude. The blue and pale-blue area around the outside of the plot shows the distribution of GC, AT, and N percentages in the same bins as the inner plot. A summary of complete, fragmented, duplicated, and missing BUSCO genes found in the assembled genome from the metazoa database (odb10) is shown on the top right.

**Figure 3.**
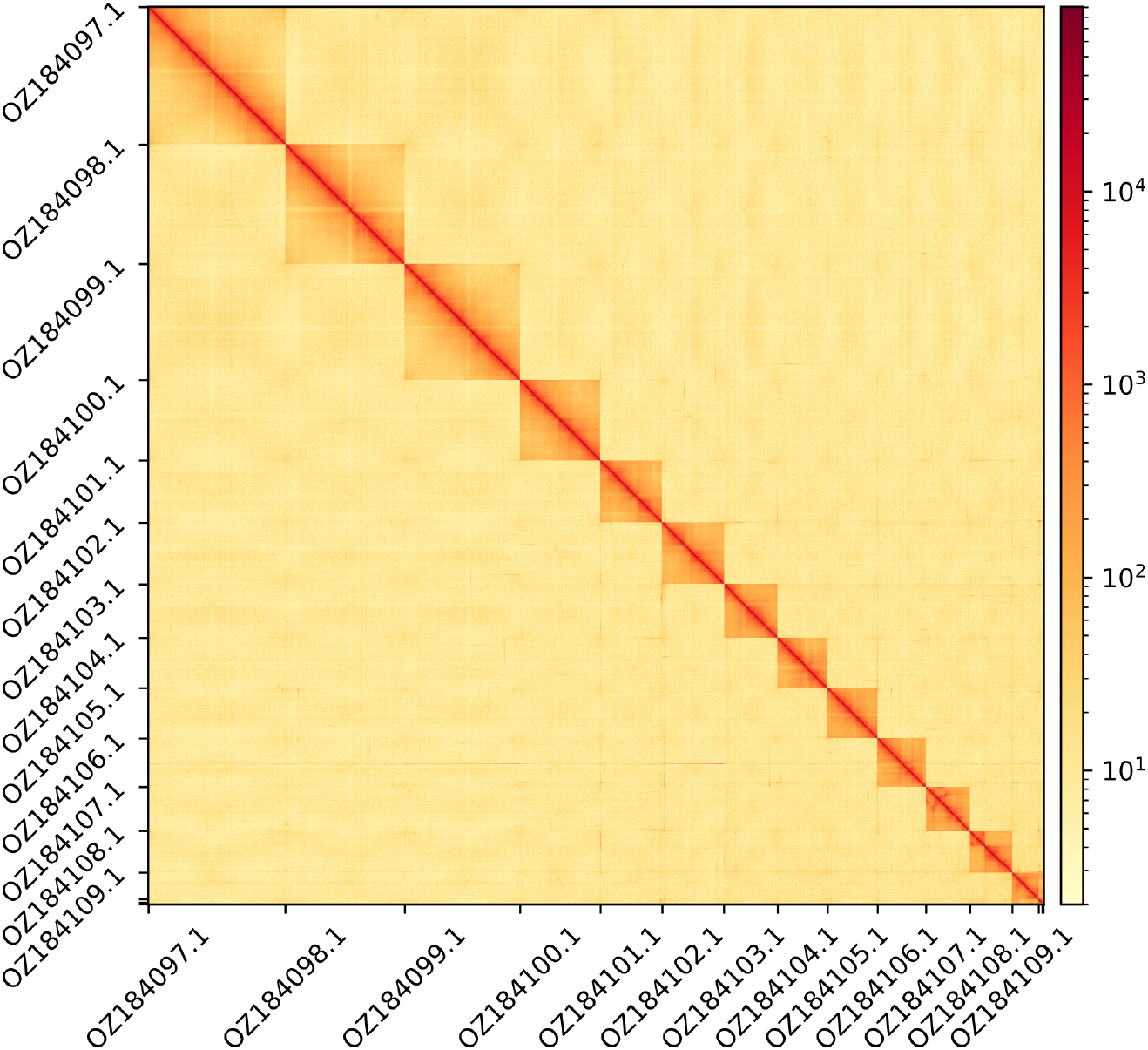
Hi-C contact map showing spatial interactions between regions of the genome. The diagonal corresponds to intra-chromosomal contacts, depicting chromosome boundaries. The frequency of contacts is shown on a logarithmic heatmap scale. Hi-C matrix bins were merged into a 50 kb bin size for plotting. The GenBank names of the 13 obtained chromosomes are shown.

## Acknowledgements

The authors want to thank Alba Enguídanos, Andrea Cabrito, Carlota Escarré, Manuel Ballesteros, Òscar Prats, and Pep Lancho for their assistance during sampling campaigns. We acknowledge the support of the Freiburg Galaxy Team: Saim Momin and Björn Grüning, Bioinformatics, University of Freiburg (Germany), funded by the German Federal Ministry of Education and Research BMBF grant 031 A538A de.NBI-RBC and the Ministry of Science, Research and the Arts Baden-Württemberg (MWK) within the framework of LIBIS/de.NBI Freiburg. We would like to acknowledge the assembly reviewer, Dominic Absolon from the Wellcome Sanger Institute.

## Author Contributions

CGC coordinated the project, MP collected the species, CGC and MP identified the species, CGC and MP sampled and preserved biological material and provided metadata, RM, TM, RO, THS, and AB provided sampling and metadata support and management, LA and MG extracted DNA, prepared libraries, and performed sequencing, FCF, JGG, and FC performed genome assembly and curation under the supervision of TSA, DDP generated the analysis and report. All authors contributed to the writing, review, and editing of this genome note and read and approved the final version.

